# Multi-modal Imaging of Disease Progression in TH-MYCN Mouse Models of Neuroblastoma

**DOI:** 10.1101/2022.03.21.484628

**Authors:** Andrew A. Badachhape, Ling Tao, Sanshuv Joshi, Zbigniew Starosolski, Laxman Devkota, Poonam Sarkar, Prajwal Bhandari, Ananth V. Annapragada, Eveline Barbieri, Ketan B. Ghaghada

## Abstract

MYCN is a major driver for neuroblastoma (NB) and the tyrosine hydroxylase (TH)-MYCN transgenic mouse model is extensively used for preclinical NB studies. However, spatio-temporal NB progression in the TH-MYCN model has not been studied, and questions remain about the value of implanted models as a surrogate for transgenic mice. In this work, we used magnetic resonance imaging (MRI) to study tumor progression and nanoparticle contrast-enhanced computed tomography (n-CECT) to assess tumor vascular architecture in TH-MYCN transgenic mice (2–7 weeks of age) and TH-MYCN+/+-derived orthotopic allograft and syngeneic mice (2–5 weeks post-tumor implantation). Tumors in TH-MYCN transgenic mice became evident in the abdominal paraspinal region at week 5. A delayed thoracic paraspinal mass became evident at week 6 and most mice succumbed by week 7. In allograft and syngeneic mice, single mass tumor growth was restricted to the peritoneal cavity. N-CECT revealed a predominantly microvascular network in TH-MYCN tumors while implanted tumors exhibited heterogeneous and tortuous vessels. N-CECT quantitative analysis demonstrated high vascularity (tumor fractional blood volume ~ 0.12) in all models. Multi-modal imaging of TH-MYCN transgenic and implanted models revealed differences in growth patterns and vascular architecture that should be considered in designing preclinical studies.

## Introduction

Neuroblastoma (NB) is the most common extracranial solid tumor in children under the age of 5 years^1,2^. Approximately 50% of NB patients present the disease with distant metastasis, most commonly to bone marrow, bone, liver and brain^1–3^ The *MYCN* oncogene plays a crucial role in tumorigenesis and angiogenesis in NB^4–7^. Amplification of *MYCN* has been correlated with an increase in tumor vascularity, high metastatic potential and poor survival in NB, and therefore is among the strongest indicators of poor prognosis^8,9^.

Over the years, a variety of mouse models have been developed to study disease initiation and progression, identify novel molecular and cellular targets, and test novel therapies, including immunotherapies^10^. The tyrosine hydroxylase (TH)-MYCN transgenic mouse model is one of the most utilized animal model of spontaneous tumor ^11,12^. growth in NB research. Mice homozygous for *MYCN* transgene selectively express *MYCN* in developing neuroblasts under the control of a rat TH promoter. Spontaneous NB tumors arise from highly proliferative progenitor cells of the sympathetic nervous system, consistent with known etiology of disease progression in pediatric NB patients^13^. While the tumor biology in TH-MYCN transgenic model has been characterized, our understanding of the spatio-temporal progression of disease has been limited in this model^14^.

Cross-sectional imaging modalities, such as magnetic resonance imaging (MRI) and computed tomography (CT) enable non-invasive, 4D (3D space + time) interrogation of disease progression^15–17^. Although *in vivo* tumor imaging in TH-MYCN NB mouse models has been attempted with MRI and nuclear imaging, 4D understanding of disease progression and tumor vascular architecture has been understudied^18–21^. In this work, we use advanced *in vivo* multi-modal imaging to study spatio-temporal disease progression in TH-MYCN transgenic mouse model of NB. Specifically, we performed high-resolution, whole-body MRI for longitudinal monitoring of NB progression. Furthermore, since *MYCN* plays a key role in tumor vascularity, contrast-enhanced CT was performed using a long circulating nanoparticle contrast agent to capture high-resolution tumor angiograms for studying age-related changes in tumor vascular architecture. In addition to studying NB progression in TH-MYCN mouse model, we applied these imaging methodologies to study and compare NB progression in orthotopic allograft and syngeneic mouse models generated using tumor cells derived from TH-MYCN transgenic mice.

## Results

Studies were performed in TH-MYCN transgenic mouse model of NB that was then used to generate orthotopic TH-MYCN-derived allograft and syngeneic models (**Figure 1**). Non-invasive monitoring of disease progression was done weekly by whole-body, non-contrast T2w-MRI. Transgenic TH-MYCN mice were imaged from two weeks to seven weeks of age (**Figure 2A**). Spontaneously growing tumors in transgenic TH-MYCN mice became evident in the abdominal paraspinal region at week 4 (~ 1.5 mm diameter). The abdominal paraspinal mass grew between the kidneys and the adrenals. As the tumors grew, the kidneys and liver were displaced by the mass and the adrenals were no longer visible. Tumor volumes computed from 3D analysis of MR images demonstrated an exponential growth phase between weeks 5 – 7 [mean volumes: 10 mm^3^ – 1500 mm^3^] (**Figure 2B**). A delayed paraspinal growth along the thoracic vertebrae became evident on MRI between weeks 6 and 7. The thoracic paraspinal mass becomes evident on MRI after tumor volume of abdominal mass exceeded 0.2 cm^3^.

**Figure 1.**
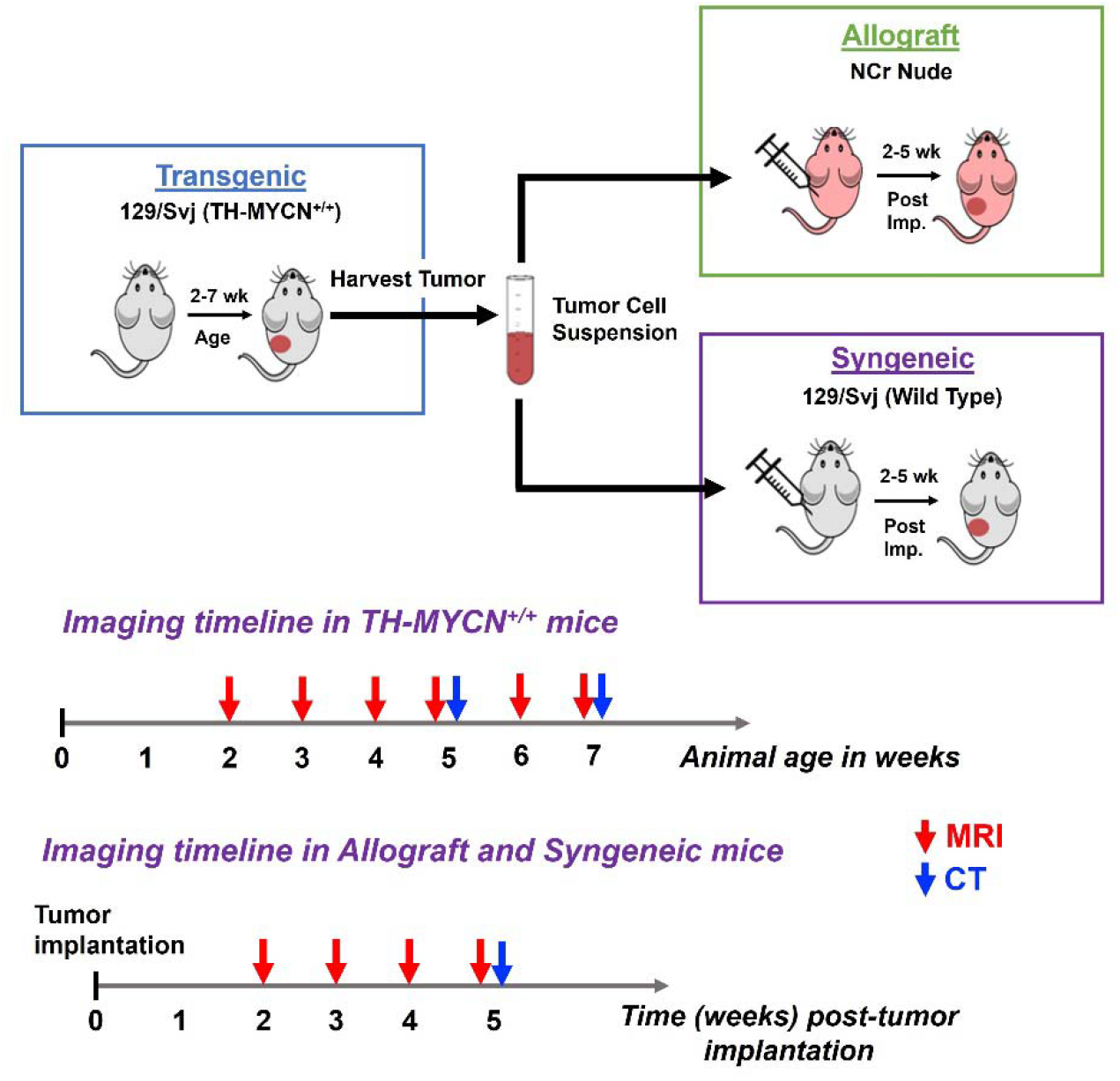
TH-MYCN transgenic and TH-MYCN-derived orthotopic allograft and syngeneic mouse models of neuroblastoma (NB). Experimental schema for generation and testing of mouse NB models. Transgenic 129×1/svj mice positive for TH-MYCN (TH-MYCN^+/+^) were identified by genotyping and monitored by MRI for tumor growth from 2 weeks of age to a terminal time point of 7 weeks of age. Tumors from TH-MYCN^+/+^ transgenic mice were harvested at terminal time point, processed into single-cell suspensions and surgically injected in the renal capsule of NCr nude mice and 129×1/svj wild type mice (TH-MYCN^-/-^) to generate TH-MYCN derived allograft and syngeneic cohorts, respectively. Allograft and syngeneic mice were monitored by MRI for tumor growth from 2 weeks post-tumor cell implantation until a terminal time point of 5 weeks post-tumor implantation.

**Figure 2.**
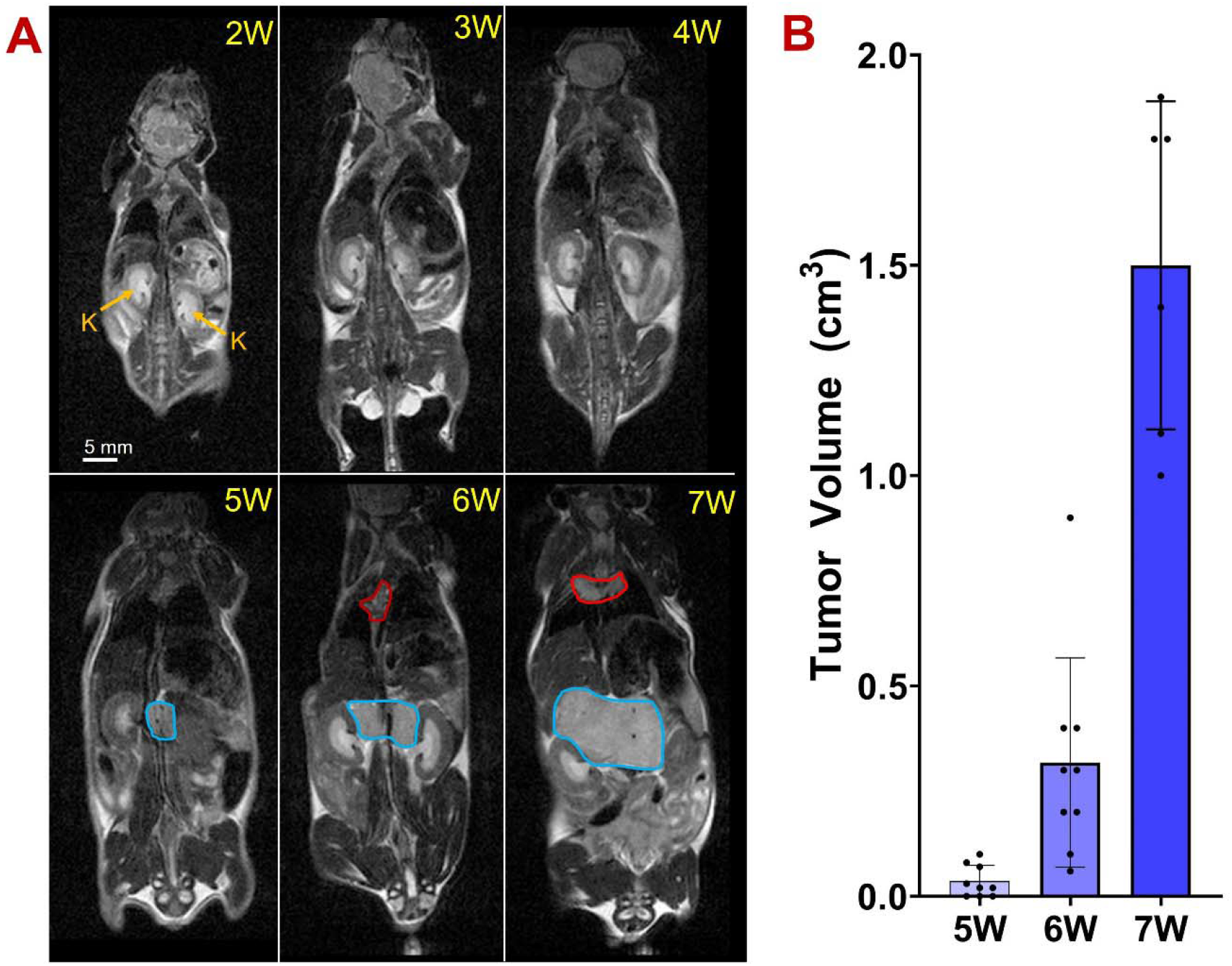
MRI of tumor progression in TH-MYCN transgenic NB mice. **(A)** Representative whole-body coronal T2-weighted MRI of tumor growth and disease progression in TH-MYCN transgenic mice. A paraspinal abdominal mass (blue contour) becomes visible on MRI at 5 weeks of age between the kidneys (K). The tumor grows rapidly and displaces adjacent organs. A delayed paraspinal thoracic mass becomes visible on MRI at 6-7 weeks of age (red contour). Scale bar represents 5 mm. **(B)** MRI-derived tumor volume (cm^3^) as a function of animal age shows an exponential growth phase between 5 and 7 weeks of age (n=9, three Tg animals were sacrificed between weeks 6 and 7 due to morbid condition).

To generate allograft and syngeneic models, tumors harvested from TH-MYCN^+/+^ mice at terminal time point (7 weeks age) were processed into single cell suspension and surgically implanted underneath left renal capsule of nude mice and 129×1/svj TH-MYCN^-/-^ wild type mice, respectively. Mice underwent non-contrast T2w-MRI from two weeks to five weeks post-tumor implantation. Unlike TH-MYCN transgenic mice, TH-MYCN-derived allograft and syngeneic mice demonstrated tumor growth restricted to the peritoneal cavity (**Figure 3A**). As disease progressed, tumors appeared to invade the left kidney (site of tumor implantation) and filled up the peritoneal space. Tumor volumes computed from analysis of MR images demonstrated a rapid growth for allograft and syngeneic tumors [mean volumes at 3 and 5 weeks post-implantation: 20 mm^3^ - 2215 mm^3^ for allograft model; 20 mm^3^ – 1730 mm^3^ for syngeneic model] (**Figure 3B**). Microscopic analysis of H&E-stained tumor sections demonstrated similarity in cellular morphology in TH-MYCN transgenic and TH-MYCN-derived allograft and syngeneic models (**Figure 3C**).

**Figure 3.**
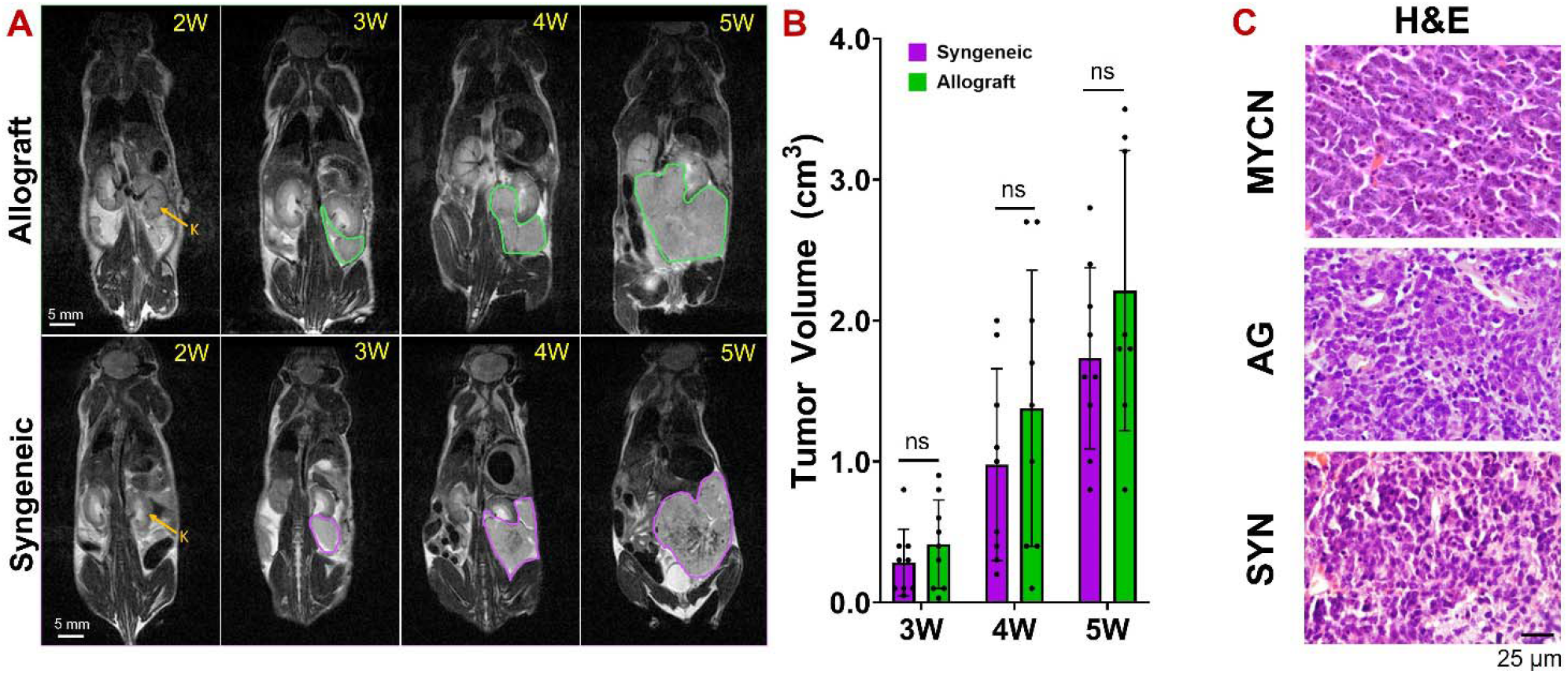
MRI of tumor progression in allograft and syngeneic NB mice. **(A)** Representative whole-body coronal T2-weighted MRI of tumor growth and disease progression in TH-MYCN derived allograft (top row) and syngeneic (bottom row) models. A solid tumor mass (red arrow) becomes visible in MRI at 3 weeks post-tumor implantation in left kidney capsule (K). Scale bar represents 5 mm. **(B)** MRI-derived tumor volume (cm^3^) shows tumor progression in TH-MYCN-derived allograft (green) and syngeneic (purple) mice until 5 weeks post-implantation (n=9); ns: not significant. One allograft mouse was euthanized between 4 and 5 week timepoints due to morbid condition. **(C)** H&E-stained images of tumors from transgenic, allograft and syngeneic mice. Scale bar indicates 25 μm.

Nanoparticle contrast-enhanced CT (n-CECT) was performed to examine tumor vasculature in TH-MYCN transgenic and implanted models. A long-circulating liposomal-iodine contrast agent was used to uniformly opacify the circulatory system and enable high-resolution CT angiogram (CTA) (35 μm isotropic spatial resolution) of tumor vasculature (**Figure 4A**). Abdominal paraspinal tumors in TH-MYCN transgenic mice demonstrated a dense network of microvasculature as evident by a predominantly homogenous signal enhancement throughout the tumor mass and lack of CT-visible large tumor-associated blood vessels. TH-MYCN mice imaged at early (5 weeks age) and late (7 weeks age) stage of disease showed similar developmental patterns of microvascular tumor architecture. The abdominal paraspinal mass surrounded the abdominal aorta as the tumor grew. The inferior vena cava was displaced and constricted by the rapidly growing mass. The thoracic paraspinal mass appeared to displace the lungs and the heart towards the caudal direction. The tumors in TH-MYCN derived implanted models exhibited markedly different vascular architecture. In contrast to tumors in transgenic mice, tumors in allograft and syngeneic mice exhibited a heterogenous tumor vascular architecture. Small (55 – 85 μm) and medium-sized (180 – 475 μm) tortuous blood vessels were seen in the core and intermediate regions of tumor. Large and tortuous superficial vessels, primarily draining venous structures, were evident in the tumor periphery (**Figure 4A**).

**Figure 4.**
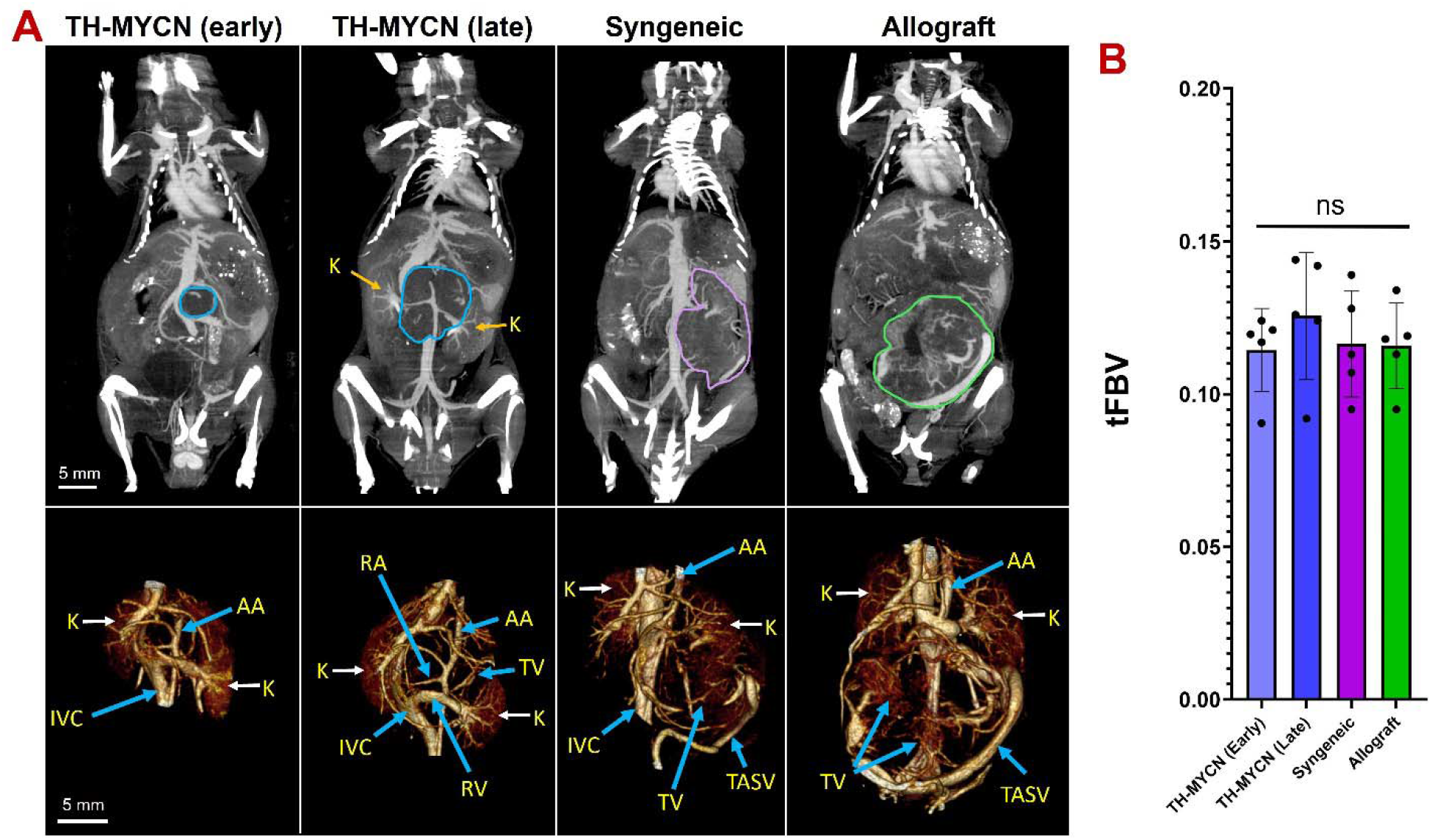
Nanoparticle contrast-enhanced computed tomography (n-CECT) imaging of tumor vasculature. **(A)** *<u>Top panel:</u>* Representative thick slab maximum intensity projection (MIP) coronal CT images depicting tumor (contours) and the vascular system; *<u>Bottom panel:</u>* 3D volume rendering of segmented tumor shows intratumoral and surrounding vasculature (*bottom panel*). Data is presented for TH-MYCN transgenic mice (5 week old “early” and 7 week old “late”), TH-MYCN-derived syngeneic and allograft mice. Scale bars represent 5 mm for both MIP images and volume renderings. K: kidney, AA: abdominal aorta, IVC: inferior vena cava. **(B)** Tumor fractional blood volume (tFBV) estimates are shown for early- and late-stage TH-MYCN transgenic (blue) and TH-MYCN-derived syngeneic (purple) and allograft (green) models. tFBV estimates were not significantly (ns) different between groups.

CT-derived tumor fractional blood volume (*tFBV*) was computed from analysis of pre-contrast and nanoparticle contrast-enhanced CT (n-CECT) images (**Figure 4B**). Despite differences in tumor vascular architecture, TH-MYCN transgenic and implanted models exhibited similar tFBV [0.12 ± 0.02 in transgenic; 0.12 ± 0.01 in allograft; 0.12 ± 0.01 in syngeneic]. Registration and fusion of MR and CT images enabled better understanding and 3D visualization of tumor development and disease progression with respect to the skeletal and blood circulatory system in TH-MYCN transgenic model (**Figure 5**). The paraspinal abdominal mass grew encompassing the abdominal aorta and its renal branches. No constriction of the aorta was observed on CT angiogram. However, the kidneys, intestines and the inferior vena cava were displaced due to the rapidly growing mass. The paraspinal thoracic mass pushed the heart towards the sternum, indicating an aggressive tumor phenotype in the TH-MYCN transgenic model.

**Figure 5.**
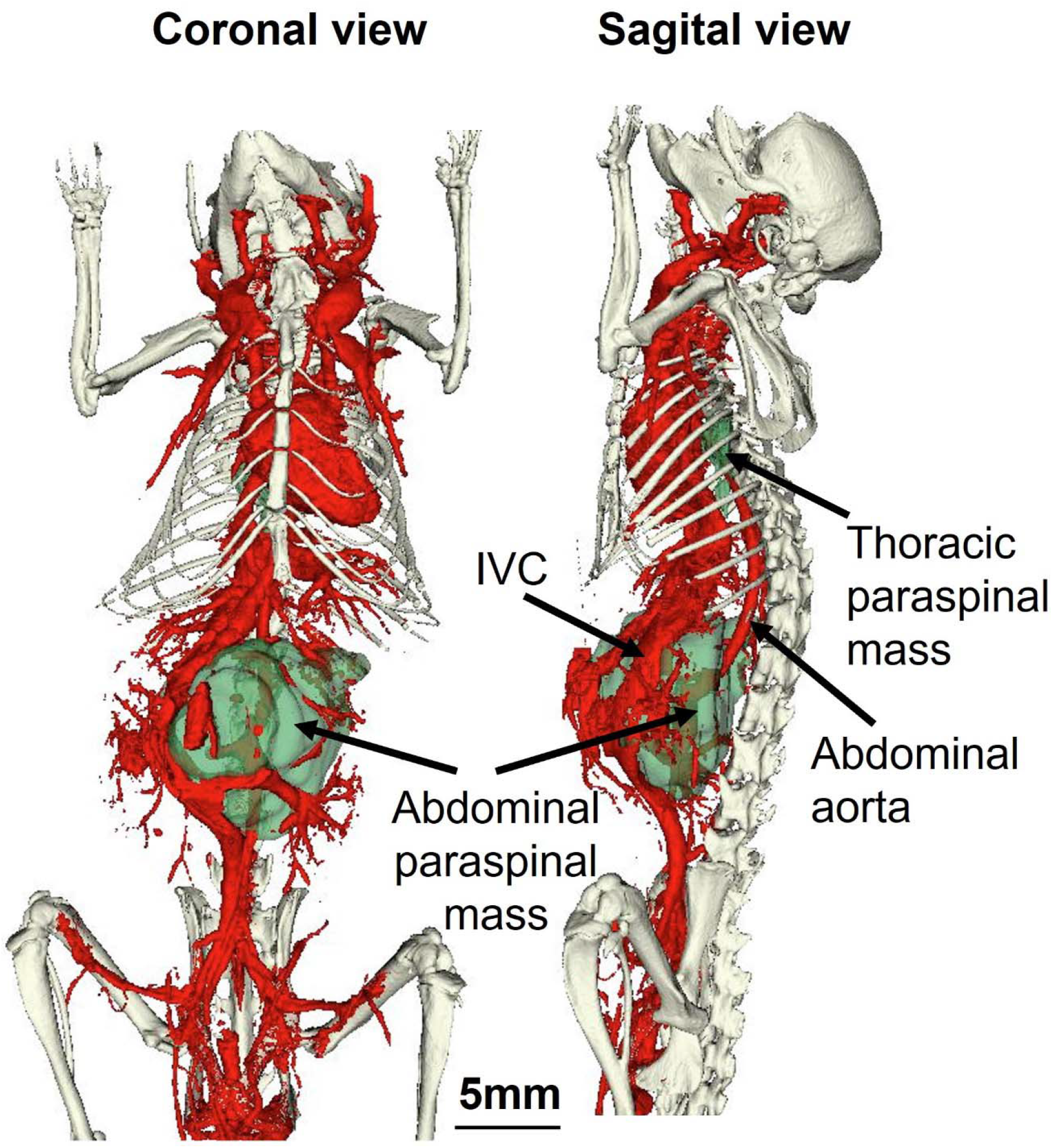
3D volume rendering of tumor masses in TH-MYCN transgenic model. 3D volume rendered whole-body coronal and sagittal views show abdominal and thoracic tumor masses (green) relative to blood circulatory system (red) and skeletal system (white). The 3D volume rendering was generated by fusing MR and CT (pre-contrast and post-contrast) datasets. Note peritoneum mass encompassing the abdominal aorta.

Pre-contrast CT images demonstrated the presence of high-attenuating focal spots, indicative of tumor calcification, in both early- and late-stage transgenic TH-MYCN animals (**Figure 6**). Analysis of pre-contrast and n-CECT images revealed the presence of calcified spots adjacent to vascular structures. Allograft and syngeneic tumors did not show evidence of calcification on CT imaging.

**Figure 6.**
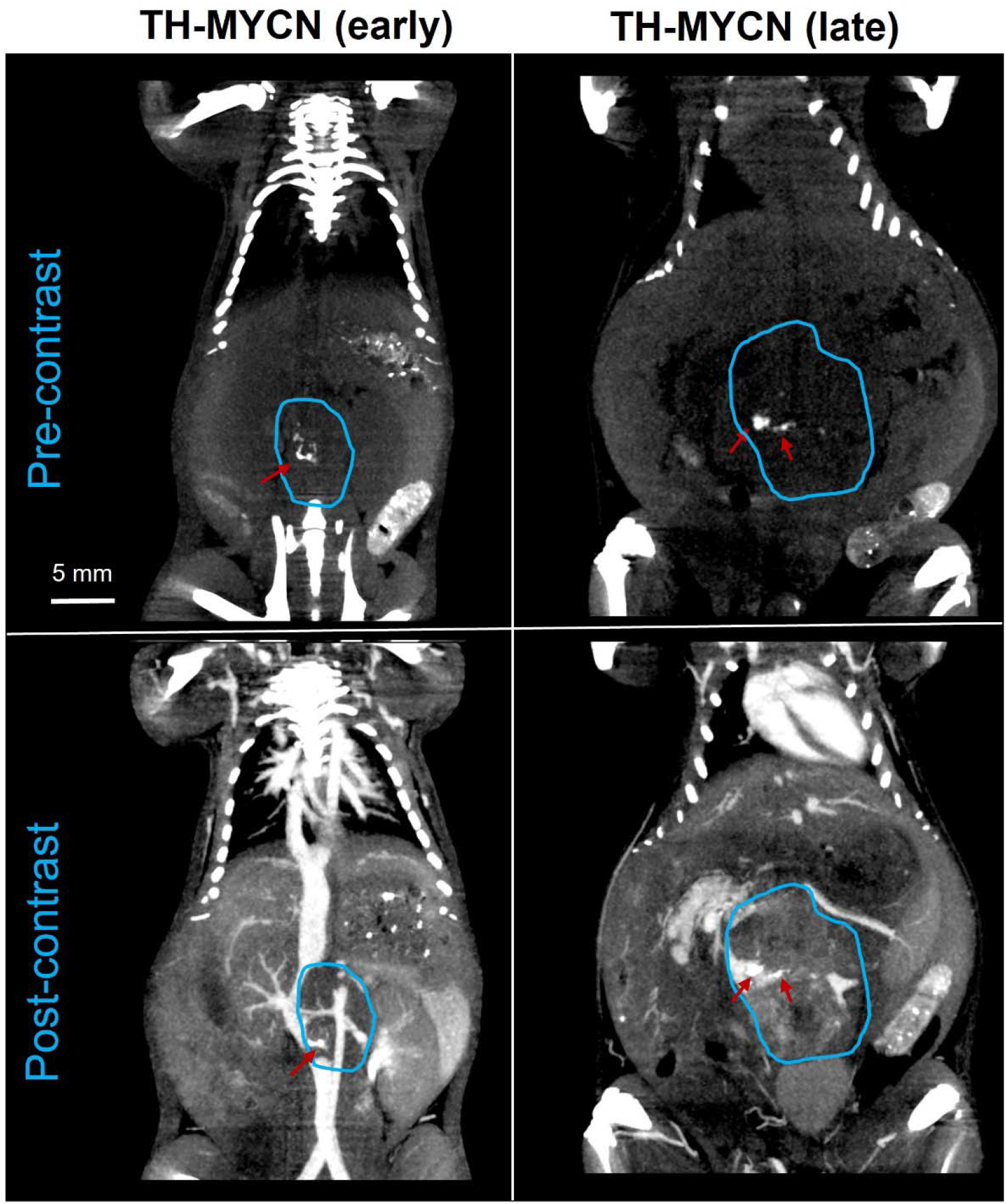
Computed tomography (CT) imaging of tumor calcification in TH-MYCN transgenic mice. Pre-contrast CT images for early and late-stage TH-MYCN animals demonstrate high attenuating regions (red arrows), indicative of tumor calcifications. Post-contrast CT images show presence of high attenuating calcified regions adjacent to tumor blood vessels.

## Discussion

Transgenic models of spontaneous tumor formation are indispensable in studying disease initiation and progression, elucidation of druggable targets and preclinical testing of novel therapies. The TH-MYCN transgenic mouse model of neuroblastoma (NB) recapitulates several aspects of disease biology seen in high-risk pediatric NB patients and therefore is commonly used to examine novel mechanisms of tumor biology. Although the transgenic model has been used for drug studies, manifestation of disease at a very young age, difficulties in breeding (~ 10% of pups are *MYCN*^+/+^) and challenges with rapid generation of large cohort of age-matched transgenic mice has attracted interest in studying alternative implantable mouse models. In this work, we used multi-modal imaging for 4D (3D imaging + time) interrogation of tumor growth and tumor vasculature in TH-MYCN transgenic and TH-MYCN derived allograft and syngeneic mouse models of NB. *In vivo* cross-sectional imaging using magnetic resonance imaging (MRI) and computed tomography (CT) enables non-invasive characterization of mouse models of solid tumors^22–24^. Furthermore, *in vivo* imaging is valuable in pre-clinical studies for non-invasive monitoring of treatment response to novel therapies^25–27^.

Longitudinal MRI revealed an aggressive disease progression in TH-MYCN transgenic NB model with spontaneous tumor originating in the paraspinal abdominal region and a delayed growth occurring along the paraspinal thoracic tract. Transgenic animals generally succumbed to the disease between 7-8 weeks of age due to the aggressive growth of masses along the paraspinal region and their increased pressure on adjacent vital organs (kidneys, heart). The exponential tumor growth and early age mortality could potentially present challenges for use of this model in therapeutic studies that incorporate a long treatment regimen. In comparison, orthotopic allograft and syngeneic renally-implanted models demonstrated single-mass tumor growth in peritoneal cavity. Tumors in implanted models did not grow as rapidly as tumors in TH-MYCN transgenic model. Further, for use in long-term therapy studies, reduction is tumor growth rate can be achieved by implanting lower number of tumor cells.

Analysis of high-resolution CT angiograms acquired using long circulating blood-pool nanoparticle contrast agent revealed a predominantly microvascular network in TH-MYCN transgenic mice. The observation of microvascular network differs from heterogenous tumor vasculature that we have previously demonstrated using n-CECT imaging in the SV-40 transgenic mouse model of NB^28^. In comparison, tumors in TH-MYCN derived allograft and syngeneic models exhibited irregular and heterogeneous vasculature including the presence of large and tortuous superficial vessels. The presence of heterogenous vasculature is consistent with our previous work in orthotopic xenograft mouse models of human NB^29^. Despite observed differences in tumor vascular architecture, all three models showed similar tumor fractional blood volume. Although not studied here, it would be interesting to evaluate and compare the ‘leakiness’ of tumor vasculature in TH-MYCN transgenic and TH-MYCN-derived implanted models.

Non-contrast CT imaging demonstrated the presence of high attenuating intratumoral features, indicative of calcifications, in TH-MYCN transgenic mice. Analysis of CT angiograms obtained using nanoparticle blood-pool contrast agent demonstrated the presence of calcification spots adjacent to blood vessels. The presence of calcifications is a key characteristic of clinical NB on CT imaging^30,31^. In comparison, TH-MYCN derived allograft and syngeneic mice did not show evidence of tumor calcifications.

The fusion of CT and MR dataset facilitated 3D understanding of tumor growth with respect to the blood circulatory and skeleton system. The paraspinal abdominal mass grew in between the kidneys encompassing the abdominal aorta and its renal branches. As the tumor grew, it pushed aside the kidneys, intestines, and the inferior vena cava. A unique characteristic of the TH-MYCN transgenic model seen in MRI and CT was the growth of paraspinal mass in the thoracic cavity at delayed time point. The paraspinal location and the exponential growth of these malignant masses in critical body regions demonstrates a highly aggressive phenotype of the TH-MYCN transgenic NB model.

In conclusion, our work describes a multi-modal imaging approach for 4D (3D imaging + time) interrogation of tumor development in TH-MYCN transgenic and TH-MYCN-derived allograft and syngeneic mouse models of NB using multi-modal imaging. This work is expected to further enhance our understanding of these tumor models and help guide future therapeutic studies for effective treatment of NB.

## Materials and Methods

### Animal Models

Studies were performed in three mouse models of NB: transgenic TH-MYCN model and two implanted TH-MYCN-derived models (allograft and syngeneic) (**Figure 1**). Homozygous transgenic 129×1/svj mice positive for TH-MYCN (TH-MYCN^+/+^) were identified by genotyping. Briefly, genomic DNA was isolated from mouse tails using DNeasy Blood and Tissue Kit (Qiagen, #69506). PCR was carried out using Econo-Tag Plus Green 2x master mix and the following primers: Chr18F1, ACTAATTCTCCTCTCTCTGCCAGTATTTGC; Chr18R2, TGCCTTATCCAAAATATAAATGCCCAGCAG; OUT1, TTGG-CACACACAAATGTATATACACAATGG. PCR condition: 93°C 3 min (1 cycle); 93°C 20 s, 56°C 20 s, 65°C 1 min 30 s (41 cycles); 72°C 7 min (1 cycle); 4°C ^32^. Both male and female transgenic mice were tested in this experiment while only female mice were used for the generation of implanted allograft and syngeneic cohorts. Disease progression in TH-MYCN transgenic mice was monitored by longitudinal MR imaging, starting at two weeks of age to a terminal time point of seven weeks of age.

For generation of TH-MYCN-derived implanted models, tumors from transgenic mice were harvested at terminal time point. Tumors were processed into single cell suspensions and injected into 2-month-old female NCr nude mice and female 129×1/svj wild type mice to generate orthotopic TH-MYCN-derived allograft and syngeneic mouse models, respectively. For both implanted models, 2.5×10^6^ tumor cells in 0.1 mL PBS were surgically injected underneath the left renal capsule using a 1ml 27gx1/2” needle^33^. Disease progression in TH-MYCN-derived allograft and syngeneic mice was monitored by longitudinal MR imaging, starting at two weeks post-tumor implantation to a terminal time point of five weeks post-tumor implantation. N=9 animals were studied in each of the three models for a total of 27 mice. Five animals in each of group also underwent nanoparticle contrast-enhanced computed tomography (n-CECT) to study tumor vascular architecture. Animals were euthanized at the terminal imaging timepoint or were sacrificed if sufficiently moribund. This was the case with three TH-MYCN mice between six and seven weeks of age and one allograft mouse that was between four and five weeks post-implantation. Mice were euthanized under isoflurane anesthesia by cervical dislocation as the primary method and then bilateral thoracic opening as the secondary method. All animal studies were performed under a protocol approved by the Institutional Animal Care and Use Committee of the Baylor College of Medicine (Protocol AN7089). All studies were in accordance with NC3RS-ARRIVE guidelines. All methods were carried out in accordance with relevant guidelines and regulations.

### Magnetic Resonance Imaging (MRI)

All animals underwent weekly whole-body MRI for monitoring of disease progression. Imaging was performed on a 1T permanent MRI scanner (M2 system, Aspect Imaging, Shoham, Israel), incorporating a 35 mm transmit-receive RF volume coil. Animals were sedated using 3% isoflurane, placed on the MRI animal bed, and then maintained at 1-2% isoflurane delivered using a nose cone setup. Respiration rate was monitored by a pneumatically controlled pressure pad placed underneath the abdominal region of the animals.

Anatomical T2-weighted (T2w) scans were acquired using a fast spin echo (FSE) sequence. Scan parameters for T2w-FSE scans were: echo time (TE) = 80 ms, repetition time (TR) = 6816 ms, slice thickness = 0.8 mm, field of view = 80 mm x 80 mm, number of slices = 33, matrix = 256 x 250, acquisition plane = coronal; in-plane resolution = 312.5 x 320 μm^2^, number of excitations = 2, echo train length = 2, scan time ~ 6 min^28,34^.

### Computed Tomography (CT) imaging

Animals underwent non-contrast, baseline CT followed by nanoparticle contrast-enhanced CT (n-CECT) for imaging of tumor vasculature and quantification of tumor fractional blood volume (tFBV). N-CECT was performed after intravenous administration of a long circulating, liposomal-iodine blood pool contrast agent (1.65 mg I/g body weight) ^28,29,35,36^. Transgenic, syngeneic, and allograft mice (n=5 for each group) were scanned on a small animal micro-CT system (Siemens Inveon) at their terminal tumor age point (7 weeks age for transgenic animals and 5 weeks post-implantation for syngeneic and allograft mice). Animals were sedated with 3% isoflurane, positioned on micro-CT bed, and then maintained under anesthesia throughout imaging session using 1-2% isoflurane delivered via nose cone. A pneumatically controlled pressure pad was placed underneath the animal’s abdominal region to monitor respiration rate during the CT imaging session.

Scan parameters for CT image acquisition were: 50 kVp, 0.5 mA, 850 ms X-ray exposure, 540 projections, 35 μm isotropic spatial resolution, scan time ~ 10 minutes. The acquired X-ray projection images were reconstructed into 3D datasets using a filtered back-projection reconstruction algorithm. All datasets were Hounsfield Unit (HU) calibrated for image analysis.

### Image analysis

Post-processing and analysis of images was performed in Osirix (version 5.8.5, 64-bit; Pixmeo SARL, Geneva, Switzerland), ImageJ (version 1.52; National Institutes of Health, USA), and MATLAB (version 2015a, MathWorks, Natick, MA) and 3D Slicer (version 4.10). Regions of interest (ROIs) were manually drawn on T2w-MR images for delineation of tumor margins and computation of MRI-derived tumor volumes.

CT-derived tumor fractional blood volume (*tFBV*) was calculated as the ratio of CT signal enhancement in tumor to blood between pre-contrast and post-contrast scans (Equation 1)^27,37^.

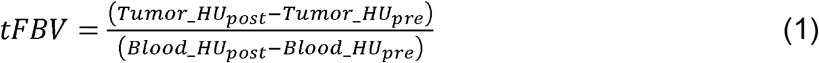

ROIs drawn in inferior vena cava (IVC) were used for blood analysis.

Total blood vessel volume was found through estimating vascular compartment volume by multiplying FBV by the tumor volumes for each individual subject.

Manual registration, segmentation, and 3D volume rendering of CT (pre-contrast), CT (post-contrast) and MR (data fusing) were performed with Slicer3d (version 4.10) software. The registration was done with a rigid transformation with 6 degrees of freedom which allowed alignment of the pelvis and chest on both modalities. The CT pre-contrast and CT post-contrast datasets were collected on the same scanner and did not require additional registration. The mice skeletal system (white) and blood circulation (red) were segmented from CT pre-contrast and CT post-contrast datasets, respectively, while the tumor mass (green) was segmented from MR.

### Histology and Immunohistochemistry

Animals were euthanized after final imaging session and tumors harvested for post-mortem analysis. Tumor tissues were weighed, fixed in 4% paraformaldehyde and embedded in paraffin. Paraffin-embedded tumor tissue blocks were sectioned and stained with Hematoxylin and Eosin (H&E). Sections were reviewed by a board-certified pathologist for microscopic evaluation of tumor’s cellular morphology.

### Statistical analysis

The non-parametric Wilcoxon rank sum test was used for statistical analysis of tumor volumes and tFBV estimates between transgenic, allograft, and syngeneic animals. GraphPad Prism 9 was used for generation of figure charts.

## Acknowledgements

The authors acknowledge the contributions of Dr. Igor Stupin who passed away after completion of this work. The authors acknowledge the Texas Children’s Hospital Small Animal Imaging Facility (SAIF) for micro-CT imaging and the Texas Children’s Hospital Pathology Core for histology. The authors acknowledge the following funding support: Gillson-Longenbaugh Foundation, NIH (NCI/NIDCR 1U01DE028233-01), St. Baldrick’s Research Grant (Grant ID:714511)

## Author Contributions

Guarantor of integrity of entire study: K.B.G.; study concepts/study design or data acquisition or data analysis/interpretation: all authors; manuscript drafting or manuscript revision for important intellectual content: all authors; approval of final version of submitted manuscript: all authors; agree to ensure any questions related to the work are appropriately resolved: all authors; literature research: A.A.B, L.T., A.V.A, K.B.G., E.B.; experimental studies: A.A.B., L.T., L.D., P.S.; statistical analysis: A.A.B., K.B.G.; manuscript editing, all authors.

## Data Availability

Data supporting the findings of this study are available from the corresponding author on request.

## References

1. Park, J. R., Eggert, A. & Caron, H. Neuroblastoma: Biology, Prognosis, and Treatment. Hematol. Oncol. Clin. North Am. 24, 65–86 (2010).

2. Nguyen, R. & Dyer, M. A. Neuroblastoma. in Neuroblastoma: Molecular Mechanisms and Therapeutic Interventions 43–61 (2019). doi:10.1016/B978-0-12-812005-7.00003-5.

3. Nakagawara, A. et al. Neuroblastoma. Jpn. J. Clin. Oncol. 48, 214–241 (2018).

4. Otte, J., Dyberg, C., Pepich, A. & Johnsen, J. I. MYCN Function in Neuroblastoma Development. Front. Oncol. 10, (2021).

5. Xue, C. et al. MYCN promotes neuroblastoma malignancy by establishing a regulatory circuit with transcription factor AP4. Oncotarget 7, 54937–54951 (2016).

6. Calero, R., Morchon, E., Johnsen, J. I. & Serrano, R. Sunitinib suppress neuroblastoma growth through degradation of MYCN and inhibition of angiogenesis. PLoS One 9, (2014).

7. Chanthery, Y. H. et al. Paracrine signaling through MYCN enhances tumor-vascular interactions in neuroblastoma. Sci. Transl. Med. 4, (2012).

8. Wang, H., Wang, X., Xu, L., Zhang, J. & Cao, H. Prognostic significance of MYCN related genes in pediatric neuroblastoma: A study based on TARGET and GEO datasets. BMC Pediatr. 20, (2020).

9. Schulte, J. H. et al. MYCN regulates oncogenic MicroRNAs in neuroblastoma. Int. J. Cancer 122, 699–704 (2008).

10. Ornell, K. J. & Coburn, J. M. Developing preclinical models of neuroblastoma: driving therapeutic testing. BMC Biomed. Eng. 1, (2019).

11. Chesler, L. et al. Malignant progression and blockade of angiogenesis in a murine transgenic model of neuroblastoma. Cancer Res. 67, 9435–9442 (2007).

12. Weiss, W. A., Aldape, K., Mohapatra, G., Feuerstein, B. G. & Bishop, J. M. Targeted expression of MYCN causes neuroblastoma in transgenic mice. EMBO J. 16, 2985–2995 (1997).

13. Terrile, M. et al. Mirna expression profiling of the murine TH-MYCN neuroblastoma model reveals similarities with human tumors and identifies novel candidate mirnas. PLoS One 6, (2011).

14. Rasmuson, A. et al. Tumor Development, Growth Characteristics and Spectrum of Genetic Aberrations in the TH-MYCN Mouse Model of Neuroblastoma. PLoS One 7, (2012).

15. Driehuys, B. et al. Small animal imaging with magnetic resonance microscopy. ILAR J. 53, 35–53 (2012).

16. Clark, D. P. & Badea, C. T. Micro-CT of rodents: State-of-the-art and future perspectives. Phys. Medica 30, 619–634 (2014).

17. Serkova, N. J. et al. Preclinical applications of multi-platform imaging in animal models of cancer. Cancer Res. 81, 1189–1200 (2021).

18. Almeida, G. S. et al. Pre-clinical imaging of transgenic mouse models of neuroblastoma using a dedicated 3-element solenoid coil on a clinical 3T platform. Br. J. Cancer 117, 791–800 (2017).

19. Zormpas-Petridis, K. et al. MRI imaging of the hemodynamic vasculature of neuroblastoma predicts response to antiangiogenic treatment. Cancer Res. 79, 2978–2991 (2019).

20. Jamin, Y. et al. Intrinsic susceptibility MRI identifies tumors with ALKF1174L mutation in genetically-engineered murine models of high-risk neuroblastoma. PLoS One 9, (2014).

21. Quarta, C. et al. Molecular imaging of neuroblastoma progression in TH-MYCN transgenic mice. Mol. Imaging Biol. 15, 194–202 (2013).

22. Zormpas-Petridis, K. et al. Noninvasive MRI native T1 mapping detects response to MYCN-targeted therapies in the Th-MYCN model of neuroblastoma. Cancer Res. 80, 3424–3435 (2021).

23. Woodfield, S. E. et al. MDM4 inhibition: a novel therapeutic strategy to reactivate p53 in hepatoblastoma. Sci. Rep. 11, (2021).

24. Iwakura, H. et al. Establishment of a novel neuroblastoma mouse model. Int. J. Oncol. 33, 1195–1199 (2008).

25. Woodfield, S. E. et al. A Novel Cell Line Based Orthotopic Xenograft Mouse Model That Recapitulates Human Hepatoblastoma. Sci. Rep. 7, (2017).

26. Shirinbak, S. et al. Combined immune checkpoint blockade increases CD8+CD28+PD-1+ effector T cells and provides a therapeutic strategy for patients with neuroblastoma. Oncoimmunology 10, (2021).

27. Devkota, L. et al. Detection of response to tumor microenvironment targeted cellular immunotherapy using nano-radiomics. Sci. Adv. 6, (2020).

28. Ghaghada, K., Starosolski, Z., Stupin, I., Sarkar, P. & Annapragada, A. Interrogation of evolving tumor vasculature using high-resolution CT imaging and a nanoparticle contrast agent. in SPIE 10578, Medical Imaging 2018: Biomedical Applications in Molecular, Structural, and Functional Imaging 47 (2018). doi:10.1117/12.2293607.

29. Ghaghada, K. B. et al. Heterogeneous uptake of nanoparticles in mouse models of pediatric high-risk neuroblastoma. PLoS One 11, e0165877 (2016).

30. Xu, Y., Wang, J., Peng, Y. & Zeng, J. CT characteristics of primary retroperitoneal neoplasms in children. Eur. J. Radiol. 75, 321–328 (2010).

31. Dumba, M., Jawad, N. & McHugh, K. Neuroblastoma and nephroblastoma: A radiological review. Cancer Imaging 15, (2015).

32. Haraguchi, S. & Nakagawara, A. A simple PCR method for rapid genotype analysis of the TH-MYCN transgenic mouse. PLoS One 4, (2009).

33. Patterson, D. M., Shohet, J. M. & Kim, E. S. Preclinical models of pediatric solid tumors (neuroblastoma) and their use in drug discovery. in Current Protocols in Pharmacology Chapter 14 (2011). doi:10.1002/0471141755.ph1417s52.

34. Badachhape, A. A. et al. Pre-clinical magnetic resonance imaging of retroplacental clear space throughout gestation. Placenta 77, 1–7 (2019).

35. Clark, D. P., Ghaghada, K., Moding, E. J., Kirsch, D. G. & Badea, C. T. In vivo characterization of tumor vasculature using iodine and gold nanoparticles and dual energy micro-CT. Phys. Med. Biol. 58, 1683–1704 (2013).

36. Mukundan, S. et al. A liposomal nanoscale contrast agent for preclinical CT in mice. Am. J. Roentgenol. 186, 300–307 (2006).

37. Badachhape, A. A. et al. Nanoparticle Contrast-enhanced T1-Mapping Enables Estimation of Placental Fractional Blood Volume in a Pregnant Mouse Model. Sci. Rep. 9, (2019).

